# Quorum sensing regulation by the nitrogen phosphotransferase system in *Pseudomonas aeruginosa*

**DOI:** 10.1101/2025.02.01.636002

**Authors:** Samalee Banerjee, Nicole E. Smalley, Pradtahna Saenjamsai, Anthony Fehr, Ajai A. Dandekar, Matthew T. Cabeen, Josephine R. Chandler

**Affiliations:** Department of Molecular Biosciences, University of Kansas, Lawrence, KS; Department of Microbiology, University of Washington, Seattle, WA; Department of Microbiology and Molecular Genetics, Oklahoma State University, Stillwater, Oklahoma

## Abstract

In the opportunistic pathogen *Pseudomonas aeruginosa*, the nitrogen-related phosphotransferase system (PTS^Ntr^) influences multiple virulence behaviors. The PTS^Ntr^ is comprised of three enzymes: first PtsP, then the PtsO phosphocarrier, and the final PtsN phosphoacceptor. We previously showed that *ptsP* inactivation increases LasI-LasR quorum sensing, a system by which *P. aeruginosa* regulates genes in response to population density. LasI synthesizes a diffusible autoinducer that binds and activates the LasR receptor, which activates a feedback loop by increasing *lasI* expression. In this study, we examined the impact of the PTS^Ntr^ on quorum sensing. Disruption of *ptsP* increased the expression of some, but not all, tested quorum-controlled genes, including *lasI, phzM* (pyocyanin biosynthesis), *hcnA* (hydrogen cyanide biosynthesis), and, to a lesser extent, *rsaL* (quorum sensing regulator). Expression of these genes remained dependent on LasR and the autoinducer, whether provided endogenously or exogenously. Increased *lasI* expression in Δ*ptsP* (or Δ*ptsO*) cells was partly due to the presence of unphosphorylated PtsN, which alone was sufficient to elevate *lasI* expression. However, we observed residual increases in Δ*ptsP* or Δ*ptsO* cells even in the absence of PtsN, suggesting that PtsP and PtsO can regulate gene expression independent of PtsN. Indeed, genetically disrupting the PtsO phosphorylation site impacted gene expression in the absence of PtsN, and transcriptomic evidence suggested that PtsO and PtsN have distinct regulons. Our results expand our view of how the PTS^Ntr^ components function both within and apart from the classic phosphorylation cascade to regulate key virulence behaviors in *P. aeruginosa*.

**IMPORTANCE:** *Pseudomonas aeruginosa* often causes severe and difficult-to-treat infections. *P. aeruginosa* virulence requires the nitrogen-related phosphotransferase system (PTS^Ntr^), which comprises the phosphocarrier proteins PtsP and PtsO and the final phosphoacceptor, PtsN. The PTS^Ntr^ is known to modulate quorum sensing, but little is known about the mechanism of regulation. Here, we examined quorum sensing regulation by the PTS^Ntr^. We showed that the PTS^Ntr^ increases quorum sensing-mediated activation of certain genes through the additive effects of both PtsO and PtsN. We also used transcriptomics to determine the regulons of PtsO and PtsN and found that they are largely nonoverlapping. The results position PtsO and PtsN as independent effectors in the Nitro-PTS and shed new light on virulence regulation in this important pathogen.

## INTRODUCTION

*Pseudomonas aeruginosa* is a Gram-negative, opportunistic pathogen found in many habitats, particularly those linked with human activity (1). *P. aeruginosa* causes severe and sometimes fatal infections in people with cystic fibrosis, acute leukemia, burn wounds and organ transplants. It is also commonly contracted in healthcare settings (2). There is a significant global health burden from *P. aeruginosa* infections, which are thought to be responsible for over $700 million in health-related costs annually (3). *P. aeruginosa* infections are particularly difficult to treat due to the prevalence of multidrug-resistant strains, an arsenal of virulence factors, and its ability to adapt and survive in diverse environments (4–6).

In *P. aeruginosa,* several key virulence factors are regulated by the nitrogen-related phosphotransferase system (PTS^Ntr^)(7). This system was first described in *Escherichia coli* as important for regulating changes in metabolism in response to the available ratio of carbon and nitrogen (8). The PTS^Ntr^ regulates diverse behaviors in different bacteria; for example, in *E. coli*, the PTS^Ntr^ regulates metabolism (9) and potassium transport (10), and the *Pseudomonas putida* PTS^Ntr^ regulates toluene degradation (11) and polyhydroxyalkanoates (12). The PTS^Ntr^ is paralogous to the canonical sugar PTSs that phosphorylate and import saccharides. The first enzyme of the PTS^Ntr^ is PtsP (“enzyme I” or EI, which is analogous to the sugar-PTS EI enzymes). PtsP transfers phosphate, thought to be taken from phosphoenolpyruvate (PEP), to the second enzyme, PtsO (“NPr,” analogous to the sugar PTS histidine protein HPr). PtsO then transfers phosphate to the final enzyme PtsN (“enzyme IIA” or EIIA, analogous to EIIA of the sugar PTS). A hallmark of the PTS^Ntr^ system as compared to sugar PTSs is that the EI has a GAF domain (named after some of the proteins it is found in; c**G**MP-specific phosphodiesterases, **a**denylyl cyclases and **F**hlA) (13); GAF domains directly bind small molecule ligands and subsequently effect a response (14). In *P. aeruginosa* and other bacteria, the *ptsO* and *ptsN* genes are located downstream of the nitrogen-related sigma factor gene *rpoN,* while the *ptsP* gene is located elsewhere (15).

The *P. aeruginosa* PTS^Ntr^ is important for virulence, although its virulence effects are not well understood. Deleting the first gene, *ptsP,* attenuates *P. aeruginosa* pathogenicity in infections of mice, *Caenorhabditis elegans* and plant leaves (16, 17). In mice, *ptsP* null mutations decrease resistance to host innate immunity (18). In laboratory evolution experiments, *ptsP* mutations increase resistance to the clinically important antibiotic tobramycin (19–21) through an unknown mechanism. In addition, the PTS^Ntr^ system impacts *P. aeruginosa* biofilm formation by modulating the Pel polysaccharide (22). Studies of biofilms also suggest that PtsO acts as a “specificity factor,” to ensure that PtsN is not phosphorylated by another PTS EI, FruB (from the fructose PTS system), and that modulation of PtsN phosphorylation impacts the expression of dozens of genes, including many that are virulence associated (7).

*ptsP* disruption also activates transcription of the quorum sensing signal synthase gene *lasI* (21, 23). Quorum sensing is a population density-dependent communication system (for reviews, see refs. (24, 25)). In *P. aeruginosa,* LasI synthesizes the acyl-homoserine lactone (AHL) signal molecule *N-*(3-oxo)-dodecanoyl L-homoserine lactone (3OC12-HSL), which is detected by the signal receptor LasR. Upon binding, LasR activates transcription of dozens of genes, including *lasI,* which creates a positive feedback loop. In addition to the LasR-LasI system, there is a second AHL signal-receptor pair in *P. aeruginosa,* RhlR-RhlI, which synthesizes and responds to *N-*butanoyl L-homoserine lactone (C4-HSL). Together, these systems activate the production of virulence factors such as pyocyanin, protease, rhamnolipids, hydrogen cyanide, biofilm matrix proteins, lectin and alkaline protease, and they have been shown to be important for virulence in numerous infection models (16, 26–29).

Although a regulatory link between quorum sensing and the PTS^Ntr^ has been established (21, 23), studies to clarify the role of *ptsP* mutation and the other two PTS^Ntr^ enzymes on quorum sensing have not been done. Using transcriptional reporters, we determined that *ptsP* disruption influences only a subset of quorum sensing-regulated genes. By adding exogenous 3OC12-HSL and using a synthetically inducible *lasI* strain, we showed that *ptsP* disruption increases LasR-dependent gene activation in response to both endogenous and exogenous 3OC12-HSL, suggesting the effects of Δ*ptsP* are not due to changes in 3OC12-HSL biosynthesis. We also demonstrated independent, but additive, effects of PtsO and PtsN on *lasI* expression; the conserved phosphorylation sites of each protein were important for differential regulation. Transcriptomics studies of strains in which the PtsO enzyme phosphorylation state varied uncovered a PtsO-dependent, distinct regulon that did not overlap with that of PtsN. The results implicate PtsO and PtsN as independent outputs of the PTS^Ntr^, highlighting the complexity of this important virulence determinant in *P. aeruginosa*.

## RESULTS

### *ptsP* deletion increases expression of some, but not all, LasR-controlled genes

To better understand the role of *ptsP-*null mutations on LasR activity, we used a pP*_lasI_-gfp* reporter plasmid (30), which contains the promoter of the LasR-responsive *lasI* gene fused to a gene encoding GFP. We transformed the pP*_lasI_-gfp* plasmid into *P. aeruginosa* Δ*ptsP* and *ΔlasR* and compared fluorescence intensities over time (Fig. 1A). These deletion mutants grew identically to the wild type strain, PA14 (Fig. S1). Consistent with our prior studies, at the end of the experiment (14 h), we observed ∼3-fold higher P*_lasI_-gfp* activation in Δ*ptsP* compared with wild type (Fig. 1A, unpaired *t-*test, p<0.001); there was no activation in the absence of LasR. This difference was observed after cultures reached an OD_600_ of ∼0.6, which correlates with the initiation of stationary phase (Fig. S1). We also measured the 3OC12-HSL concentration in wild type and Δ*ptsP* cultures after 18 h growth. We found that 3OC12-HSL levels were almost 5-fold higher for the Δ*ptsP* strain compared with that of the wild type; 279 nM for Δ*ptsP* and 57 nM for the wild type (unpaired *t-*test, p < 0.0001), consistent with the observed difference in *lasI* transcription in these two strains.

**Fig. 1.**
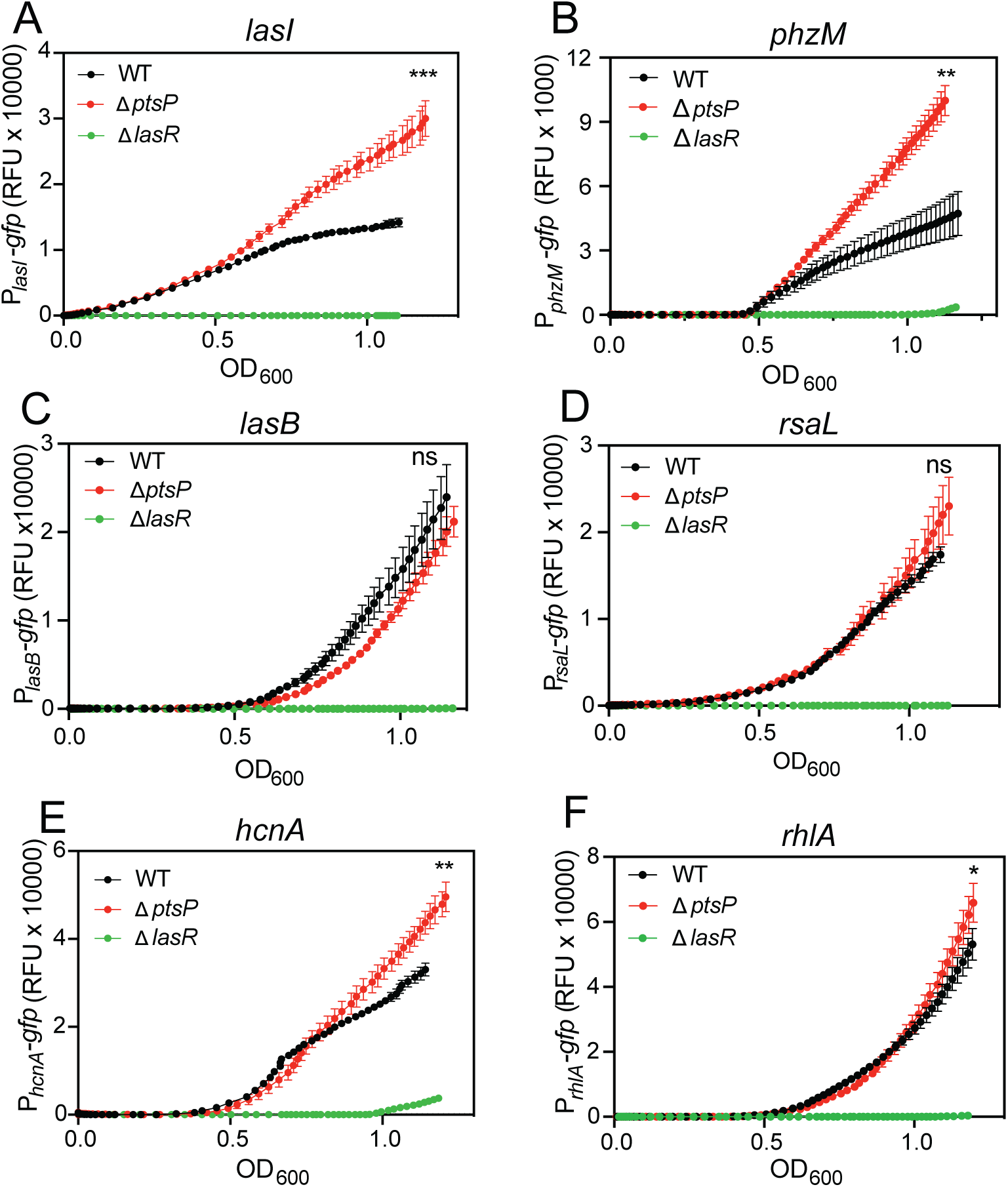
Δ*ptsP* influence on quorum sensing gene transcription. GFP fluorescence was measured in strains harboring reporters to the *lasI* (A), *phzM* (B), *lasB* (C), *rsaL* (D), *hcnA* (E), and *rhlA* (F) promoters in PA14, PA14 *ΔptsP* and PA14 Δ*lasR.* Fluorescence output was measured over a time course using a BioTek plate reader. Data points are the means of three replicates, and the error bars represent standard deviation. Significance by t-test of wild type compared with Δ*ptsP* from the final time point (OD_600_-adjusted fluorescence values); ***, p < 0.0005; **, p < 0.005; *, p < 0.05; ns, not significant.

We sought to test if *ptsP* disruption similarly regulates expression of other quorum sensing-regulated genes. We selected promoters for the following genes: *phzM*, encoding a key enzyme in phenazine biosynthesis, based on prior studies showing PTS^Ntr^ regulation of phenazine production (21, 23); *lasB* (elastase protease), *rsaL* (repressor of LasR-I system), *hcnA* (hydrogen cyanide biosynthesis), and *rhlA* (rhamnolipid surfactant)(31). This suite of genes as a group tests the activity of both RhlR and LasR quorum sensing. We engineered plasmid-based reporters of each of these gene promoters (pP*_phzM_-gfp,* pP*_lasB_-gfp,* pP*_rsaL_-gfp,* pP*_hcnA_-gfp,* and pP*_rhlA_-gfp*). As expected, the expression of each of these genes was dependent on LasR in our time course experiments (Fig. 1). Compared with wild type, in the Δ*ptsP* strain we observed higher activation of the *phzM, hcnA,* and, to a lesser degree *rsaL* reporters. These results show disrupting *ptsP* increases expression of some, but not all, quorum sensing-regulated genes.

### Δ*ptsP*-dependent *lasI* regulation requires LasR and 3OC12-HSL

We sought to gain further insight into the relationship of PtsP to *lasI* regulation. We first asked if LasR is required for Δ*ptsP-*dependent activation of *lasI* transcription. We deleted *lasR* from a Δ*ptsP* mutant and measured *lasI* expression levels using our pP*_lasI_-gfp* reporter. In the absence of *lasR,* we observed no significant difference in *lasI* activation in a Δ*ptsP* mutant compared with wild type (Fig. 2A), supporting the idea that elevated *lasI* expression in a Δ*ptsP* mutant requires LasR.

**Fig. 2.**
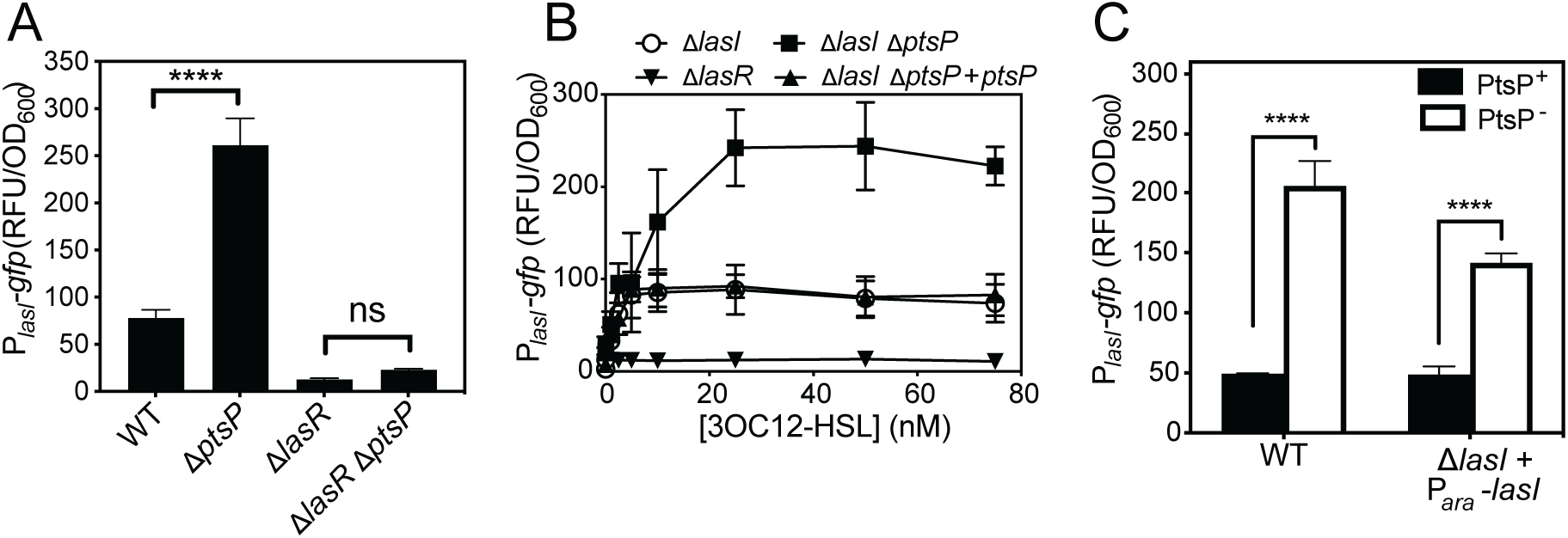
Role of LasR and exogenous 3OC12-HSL on Δ*ptsP-*dependent *lasI* activation. Growth-adjusted GFP fluorescence in strains harboring the P*_lasI_-*GFP reporter plasmid. Fluorescence was measured after 18 h growth in test-tube grown cultures. For panels B and C, strains carried a chromosomally inserted CTX-1 cassette (CTX), CTX plus *ptsP,* or CTX plus the arabinose-inducible *lasI* gene (Para-*lasI*). For (C), black bars indicate strains with the native *ptsP* gene intact and white bars indicate strains with Δ*ptsP*; 0.2% arabinose was added to all cultures. Data points are the means of three replicates, and the error bars represent standard deviation. Significance by t-test; ***, p < 0.0005; ns, not significant.

Next, we considered whether the Δ*ptsP* mutation increases biosynthesis or production of 3OC12-HSL, which could increase *lasI* expression via activation of LasR. We used two approaches to test this hypothesis. First, we deleted *lasI* from our wild type and Δ*ptsP* mutant strains and grew these strains with varying concentrations of exogenously added 3OC12-HSL (Fig. 2B). With the pP*_lasI_-gfp* reporter, the minimum 3OC12-HSL concentration we could detect a response was 1 nM. The response was saturated at concentrations of 25 nM and higher. In this concentration range, we observed a ∼3-fold increase in GFP levels in the Δ*ptsP* strain. We could restore GFP levels to the wildtype level this strain by inserting an intact copy of the *ptsP* gene into the neutral *att* site in the chromosome. We also engineered strains to constitutively express *lasI* using an arabinose-inducible promoter (P*_ara_*-*gfp*), which we inserted into the neutral *att* site in the chromosome of our Δ*lasI* and Δ*lasI-*Δ*ptsP* mutants. This approach removes *lasI* expression from LasR control (32). With these strains, we also observed a ∼3-fold increase in P*_lasI_-gfp* activation due to the Δ*ptsP* mutation. The Δ*lasI* and Δ*lasI-*Δ*ptsP* strains harboring P*_ara_*-*lasI* produced 3OC12-HSL at concentrations of 254 nM and 198 nM, respectively, which is sufficient to maximally induce the GFP reporter (Fig. 2B). From these results, we conclude that deleting *ptsP* increases LasR sensitivity to 3OC12-HSL, rather than modulating 3OC12-HSL biosynthesis.

### Deleting the PtsP GAF domain does not impact *lasI* expression

In *E. coli,* the phosphorylation state of PtsP appears to depend on the cellular ratio of nitrogen to carbon, which is detected by direct binding of glutamine and α-ketoglutarate by PtsP through the GAF domain (Fig. 3A). In *P. aeruginosa,* the GAF domain appears to be required for PtsP to transfer phosphates to PtsO and PtsN (7). We thus asked whether the PtsP GAF domain is required for its role in *lasI* regulation. Therefore, we constructed a PA14 strain in which we deleted the GAF domain from *ptsP* at its native site in the genome (*ptsP*ΔGAF). We introduced our pP*_lasI_-gfp* reporter plasmid to this *ptspΔ*GAF strain and compared *lasI* transcription activation with that of the wild type and a full *ptsP* deletion mutant. As expected, *lasI* expression increased in the Δ*ptsP* strain; however, deleting only the GAF domain did not similarly increase *lasI* expression (Fig. 3B). We also tested whether the PtsP GAF domain is required for PtsP’s role in regulating the *phzM* or *hcnA* promoters by using the pP*_phzM_-gfp* and pP*_hcnA_-gfp* reporter plasmids, respectively. As with *lasI,* we did not observe a significant role for the PtsP GAF domain in regulating *phzM* or *hcnA* expression (Fig. S2). These results show the GAF domain is dispensable for PtsP’s role in regulating *lasI, phzM* and *hcnA*.

**Fig. 3.**
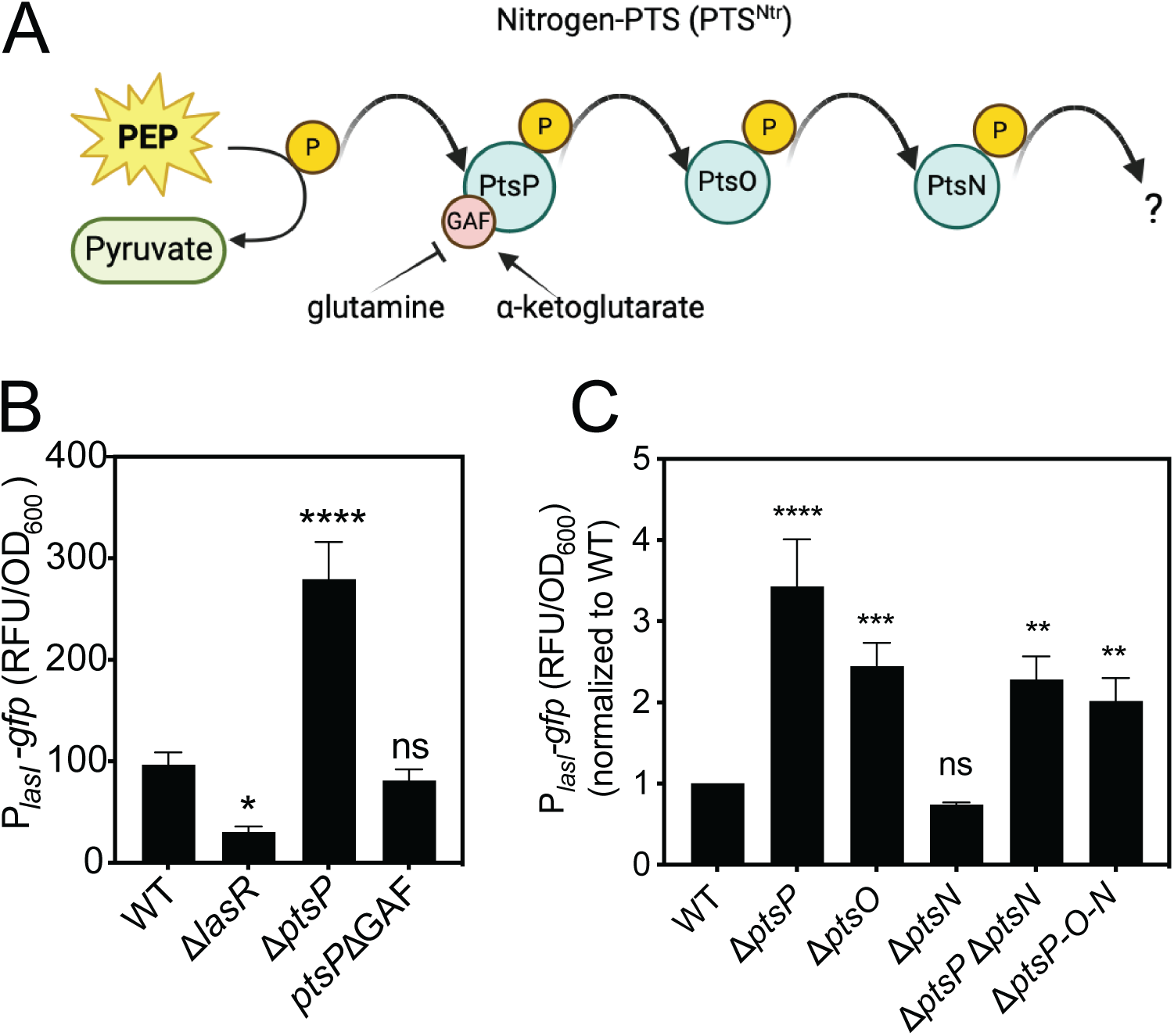
The PtsO and PtsN enzymes, but not the PtsP GAF domain, contribute to *lasI* regulation. (A) Illustration of the PTS^Ntr^ phosphorylation cascade. Phosphate is transferred by PtsP (EI Ntr) from phosphoenolpyruvate (PEP) to PtsO (Npr) and in turn to PtsN (EII Ntr), which has unknown regulatory targets. The GAF domain of PtsP is thought to regulate phosphorylation by detecting changes in the ratio of glutamine and α-ketoglutarate. (B) and (C) Transcription from the *lasI* promoter was monitored as GFP fluorescence in cells transformed with the pP*_lasI_-gfp* reporter plasmid. *ptsP*ΔGAF is an allelic replacement of *ptsP* with a deletion of the sequence encoding the GAF domain. Δ*ptsP-O-N* is the Δ*ptsP-*Δ*ptsO*-Δ*ptsN* triple mutant. Data shown are growth-adjusted fluorescence (B) or growth-adjusted fluorescence normalized to wild type (C) after 18 h growth. Data are means of at least three replicates, and error bars represent SD. Statistical significance by one-way ANOVA compared with wild type; ****, p < 0.0001; **, p < 0.005; ns, not significant.

### *lasI* is regulated by other enzymes in the PTS^Ntr^ pathway

Next, we examined the role of PtsO and PtsN, the other two enzymes in the PTS^Ntr^ phosphotransfer system (Fig. 3A), for *lasI* regulation. As a first step, we introduced the P*_lasI_-gfp* reporter plasmid to wild-type PA14 and single, double and triple deletion mutant(s) of the three PTS^Ntr^ genes *ptsP*, *ptsO* and *ptsN* (Fig. 3C). Deleting *ptsO* from the wild-type genome increased *lasI* expression to nearly the same level as the Δ*ptsP* mutant (∼2-fold for Δ*ptsO* and ∼3-fold for Δ*ptsP*) (Fig. 3C). In the Δ*ptsO* mutant, we could restore *lasI* expression to that of the wild type by introducing a functional copy of *ptsO* into the genome (Fig. S3). These results show that *lasI* is regulated, in part, by PtsO. Deleting *ptsN* from the wild-type genome did not significantly alter *lasI* expression (Fig. 3C); however, deleting *ptsN* from the Δ*ptsP* mutant decreased *lasI* expression but not to wild type levels. Thus, PtsN regulates transcription from the *lasI* promoter, but only in the absence of PtsP. In a strain in which all three of the PTS^Ntr^ enzymes are deleted (Δ*ptsP-ΔptsO-ΔptsN*), we observed an increase in *lasI* expression as compared to wild type. Together, the results support the idea that PTS^Ntr^ may have regulatory effects on *lasI* expression that are both activating (in the absence of PtsP or PtsO) and suppressing (when PtsN, or PtsO and PtsN, are absent in a Δ*ptsP* background).

### Unphosphorylated PtsN activates *lasI* expression

PtsN is unphosphorylated in the absence of PtsP (7), leading us to posit that *lasI* activation in a Δ*ptsP* mutant may be at least partially due to unphosphorylated PtsN. This model is consistent with our finding that deleting *ptsN* from a Δ*ptsP* mutant reduces *lasI* expression in this strain (Fig. 3C). To test this hypothesis, we utilized a PtsN allele that harbors a single amino acid substitution (H68A) that changes its phosphorylation site to alanine. This substitution has been shown to effectively block PtsN phosphorylation in *P. aeruginosa* (7). We moved the unmutated PtsN or the PtsN^H68A^ genes in single copy to the chromosome of the Δ*ptsN* mutant and used the pP*_lasI_-gfp* reporter plasmid to compare *lasI* expression in these strains with that of the Δ*ptsN* and wild type strains (Fig. 4A and B). Consistent with our earlier result, *lasI* expression was similar in the wild type and Δ*ptsN* mutants. In the Δ*ptsN* strain, we found that PtsN^H68A^, but not the wild-type PtsN, increased *lasI* expression by ∼3-fold; this difference was observed after cultures reached stationary phase, with no significant effects on growth (OD_600_ ∼0.6; Fig. S4). We also assessed the role of PtsN and PtsN^H68A^ in regulating *lasI* expression in the Δ*ptsP-ΔptsO-ΔptsN* mutant, where we could evaluate regulation effects in the absence of the other PTS^Ntr^ enzymes. In this genetic background, we expected *lasI* expression to be activated by ectopically expressing PtsN^H68A^, as observed when we expressed this allele in the Δ*ptsN* strain. We also expected *lasI* expression to be similarly activated with wild-type PtsN in this strain, because PtsN will be unphosphorylated in the absence of *ptsP.* Consistent with our expectations, PtsN^H68A^ increased *lasI* expression levels ∼2.4-fold compared with no PtsN alleles. However, wild-type PtsN increased levels to a lesser degree than that of the PtsN^H68A^ allele, by only ∼1.4-fold (Fig. 4C). We speculate that PtsN may be partially phosphorylated in this strain by FruB, the fructose EI enzyme, which can phosphorylate PtsN in the absence of PtsO (7). Thus we also examined the role of PtsN and PtsN^H68A^ on *lasI* expression in a Δ*ptsP ΔptsN* mutant, where PtsO is intact. In this strain, transcription from the *lasI* promoter was increased to the same level by ectopically expressing either PtsN^H68A^ or PtsN (Fig. S4). Together, these results support the conclusion that unphosphorylated PtsN activates *lasI* expression, and disrupting *ptsP* results in elevated transcription levels due to an accumulation of unphosphorylated PtsN.

**Fig. 4.**
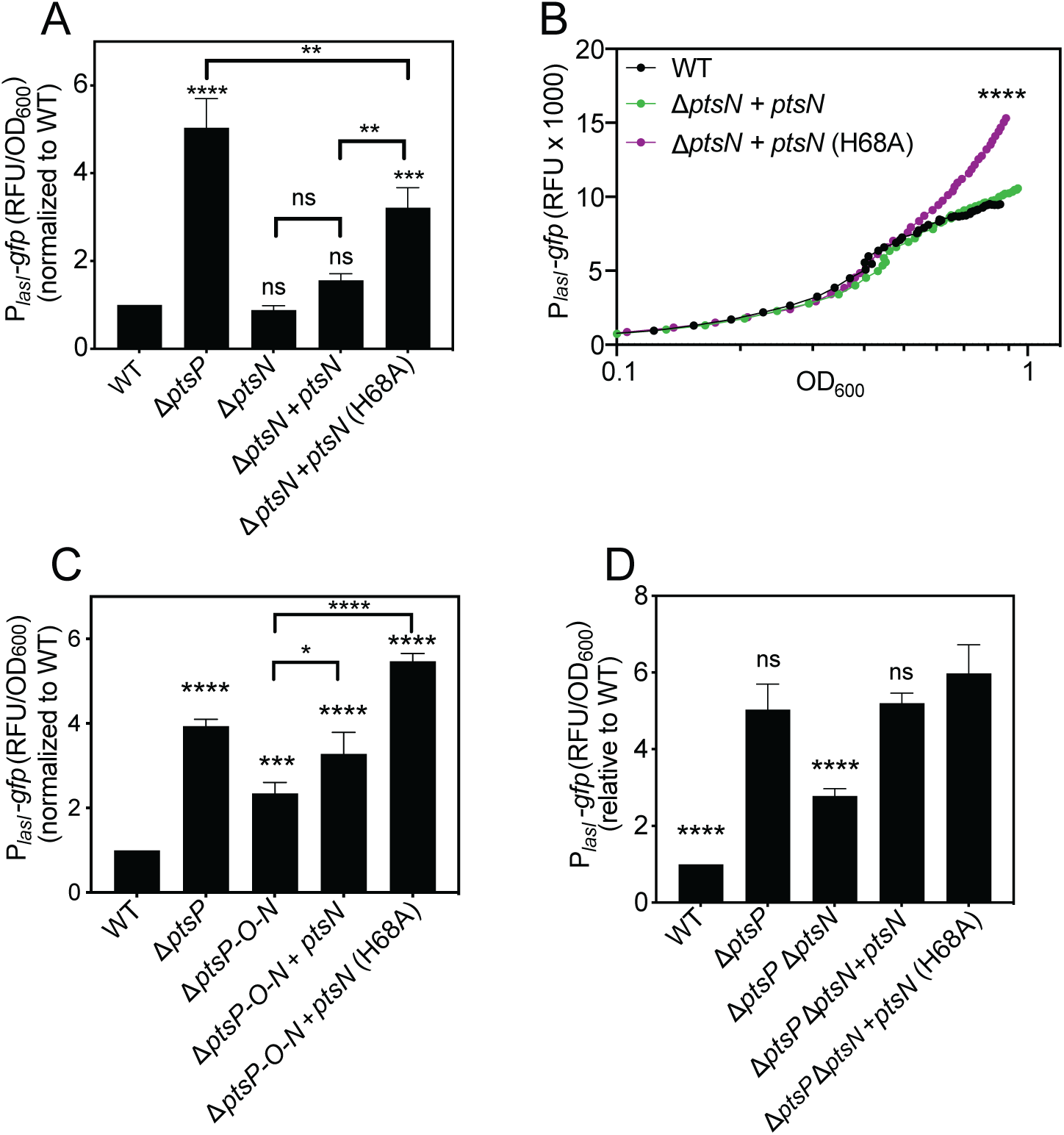
Mutating the PtsN phosphorylation site increases *lasI* expression. Transcription from the *lasI* promoter was monitored as GFP fluorescence in cells transformed with the pP*_lasI_-gfp* reporter plasmid. Panels A, C and D show growth-adjusted fluorescence normalized to wild type after 18 h of growth, and panel B shows fluorescence output measured over a time course using a BioTek plate reader. Strains carried a chromosomally inserted CTX-1 cassette (CTX), CTX plus the wild type *ptsN,* or CTX plus the H68A variant of PtsN (*ptsN* (H68A)). Data are means of at least three replicates, and error bars represent SD (error bars are too small to be seen in panel B). Statistical significance by one-way ANOVA compared to wild type (or as indicated) for panels A-C and compared to Δ*ptsP-ΔptsN* + *ptsN-*H68A for panel D; ****, p < 0.0001; ***, p<0.001; **, p < 0.005; ns, not significant. Statistical analysis of data shown in panel B were of OD-adjusted fluorescence values from the final time point.

### In the absence of PtsN, *lasI* expression is repressed by both PtsP and PtsO

In our experiments, *lasI* reporter activity was significantly lower in the Δ*ptsN* mutant than in the Δ*ptsP-ΔptsN* mutant (Fig. 3C), suggesting there is another mechanism of PTS^Ntr^ regulation that may be through PtsP or PtsO. Thus, we sought to unravel this additional regulation mechanism. We reasoned that if *lasI* is repressed by PtsP only, we should observe a PtsP-dependent reduction of *lasI* reporter activity in the absence of both PtsO and PtsN. To test this hypothesis, we moved *ptsP* in single copy into the chromosome of the Δ*ptsP-ΔptsO-ΔptsN* triple mutant and transformed these strains with the P*_lasI_*-*gfp* plasmid reporter to measure transcription from the *lasI* promoter. We found that *ptsP* had no effect on *lasI* expression in this strain (Fig. 5A), showing that PtsP alone does not modulate *lasI* expression. However, in the Δ*ptsP-ΔptsN* mutant (where *ptsO* is intact), *ptsP* significantly decreased *lasI* expression (Fig. 5A); differences in expression, but not growth, were observed after cultures reached stationary phase (OD_600_ ∼0.6; Figs. 5B and S5A). These results suggest that PtsP can repress *lasI* expression but only in the presence of PtsO. To determine whether PtsO can regulate *lasI* on its own, we compared *lasI* expression in the Δ*ptsP-ΔptsO-ΔptsN* triple mutant with that of the Δ*ptsP-ΔptsN* double mutant (where *ptsO* is intact) (Fig. 5A). However, *lasI* expression in these two strains was indistinguishable (Fig. 5A), indicating PtsO alone does not modulate *lasI* expression. From these data we conclude that *lasI* suppression in the absence of PtsN requires both PtsP and PtsO.

**Fig. 5.**
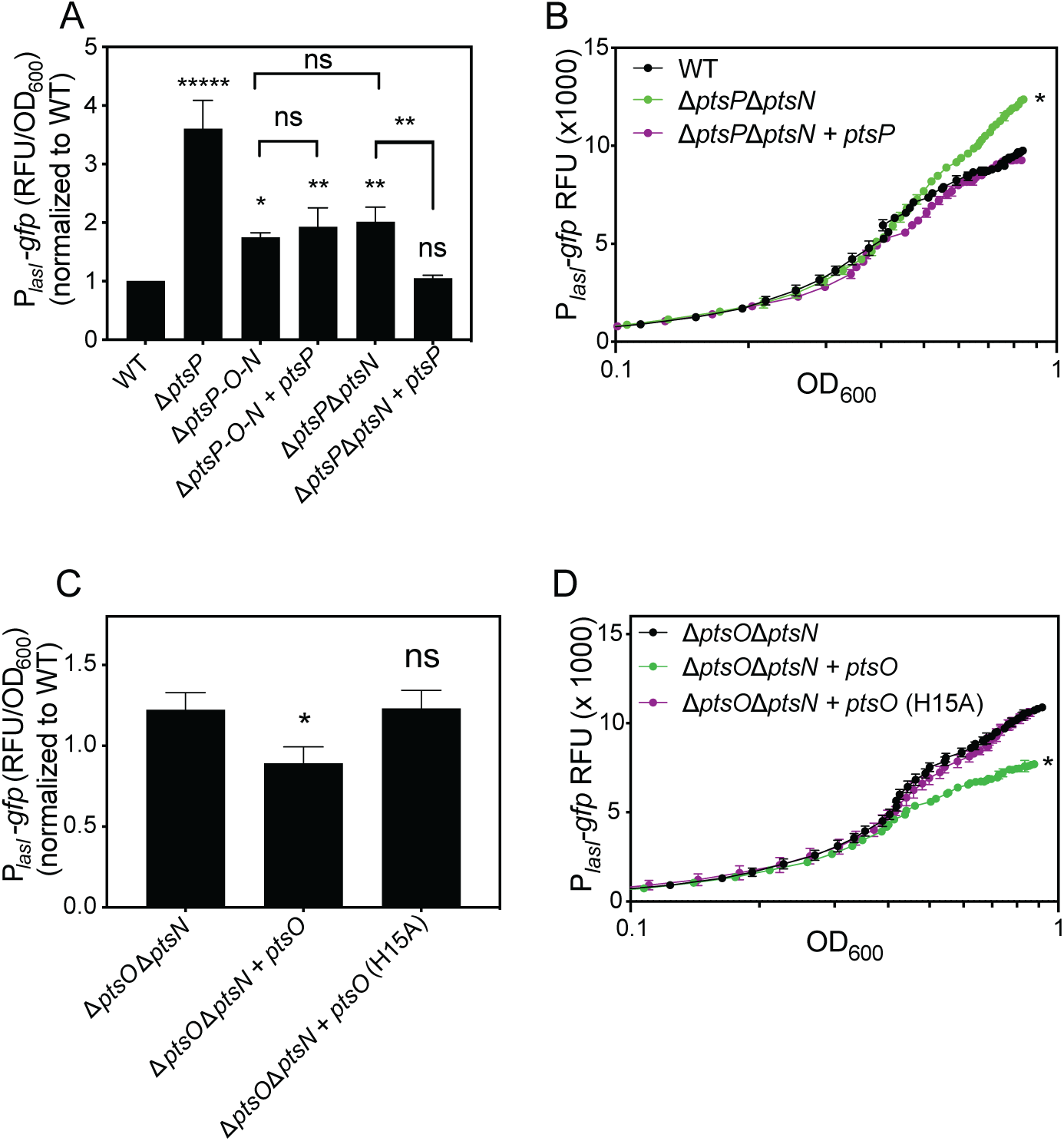
*lasI* regulation by PtsP and PtsO. Transcription from the *lasI* promoter was monitored as GFP fluorescence in cells transformed with the pP*_lasI_-gfp* reporter plasmid. (A and C) Growth-adjusted fluorescence normalized to wild type after 18 h of growth. (B and D) Fluorescence output measured over a time course using a BioTek plate reader. Strains carried a chromosomally integrated CTX-1 cassette (CTX), CTX plus the wild type *ptsO,* or CTX plus the H15A variant of PPtsO (*ptsO* (H15A)). Data are means of three replicates, and error bars represent SD (error bars are too small to be seen in panel B). Statistical significance by one-way ANOVA compared with the wild type (A and B) or Δ*ptsOΔptsN* (C and D) *, p < 0.05; ns, not significant. Statistical analysis of data shown in panel B and D were of OD-adjusted fluorescence values from the final time point.

### PtsO represses *lasI* transcription through its conserved phosphorylation site

The requirement for both PtsP and PtsO in suppressing *lasI* transcription implied that PtsO phosphorylation, which depends on PtsP, might have a regulatory role. Thus, we considered the possibility that phosphorylated PtsO represses *lasI* expression. To test this hypothesis directly, we ectopically expressed *ptsO* from the neutral *attB* site in the chromosome of the Δ*ptsO-ΔptsN* mutant (where PtsP is present). In this mutant, introducing *ptsO* caused a small but significant decrease in *lasI* expression (Fig. 5C), which correlated with stationary phase in time-course experiments (Fig. 5D and S5B). Next, we constructed and tested a *ptsO* allele in which the conserved phosphorylation site (His15) is substituted with an alanine. PtsO has a high degree of conservation across many bacterial species, and the His15 phosphorylation site is universally conserved across this family (33). Mutation of this residue to alanine blocks phosphotransfer to PtsN and disrupts PTS^Ntr^-controlled phenotypes in the related species *Pseudomonas putida* (34). To test the hypothesis that the PtsO His15 residue is important for it to repress *lasI*, we moved the PtsO^H15A^ gene into the chromosome of the Δ*ptsO-ΔptsN* strain and assessed activation from the *lasI* promoter using the P*_lasI_*_-_*gfp* reporter plasmid. We did not observe any decrease in transcription from *the* lasI promoter in the strain with the PtsO^H15A^ allele as we did for PtsO (Fig. 5C and D). There was also no significant effect on growth or PtsO levels in whole-cell lysates that could explain this result (Fig. S5). Although there are other potential explanations, such as misfolding of the PtsO^H15A^ protein, these results are consistent with the idea that PtsO represses *lasI* expression due to phosphorylation at its His15 residue.

In our experiments, the ability of PtsO to repress *lasI* transcription did not require PtsN, suggesting that PtsO repression is independent of its role in phosphotransfer to PtsN, the terminal acceptor. There are two other enzymes in *P. aeruginosa* with terminal phosphoacceptor activity analogous to that of PtsN; these are the carbohydrate PTS enzymes FruB and NagF. Thus, we sought to test the hypothesis that FruB or NagF could serve as alternative phosphor-recipients from PtsO. We expected that if FruB or NagF are important for PtsO to regulate *lasI,* then deleting *fruB* or *nagF* from the wild-type genome would increase *lasI* expression similar to deleting *ptsO*. However, deleting *fruB* and *nagF* had no measurable effects on *lasI* expression (Fig. S6). These results support the idea that PtsO represses *lasI* expression independently of other PTS enzymes.

### PtsO regulates a distinct subset of genes

PtsO is not known to modulate gene transcription independent of its role in phosphotransfer to PtsN in *P. aeruginosa.* We hypothesized that it might and sought to determine if PtsO similarly regulates genes by a PtsN-independent mechanism. First, we examined *phzM* and *hcnA* because expression from these promotors was affected by the Δ*ptsP* mutation (Fig. 1). For this experiment, we introduced the P*_phzM-_gfp* and P*_hcnA_-gfp* reporter plasmids to the Δ*ptsO-ΔptsN* mutant with ectopically expressed PtsO, PtsO^H15A^, or neither, and compared fluorescence intensities over a time course. We observed a small but significant reduction of *phzM* reporter activity due to expression of PtsO that was not observed with PtsO^H15A^ (Fig. 6A) and not explained by differences in growth (Fig. S7A); however, we observed no PtsO-dependent differences in *hcnA* reporter activation (Fig. S8). These results support the idea that phosphorylated PtsO decreases transcription from the *phzM* promoter, but not the *hcnA* promoter. We also assessed whether *phzM* transcription is regulated by PtsN. To test this, we introduced the P_phzM_-gfp reporter to the Δ*ptsN* strain with ectopically expressed PtsN, PtsN^H68A^, or neither. We found that PtsN^H68A^, but not PtsN, increased *phzM* expression >3-fold compared with (Fig. S7B). Thus both PtsO and PtsN modulate transcription from the *phzM* promoter in a manner that is similar to that of *lasI*.

**Fig. 6.**
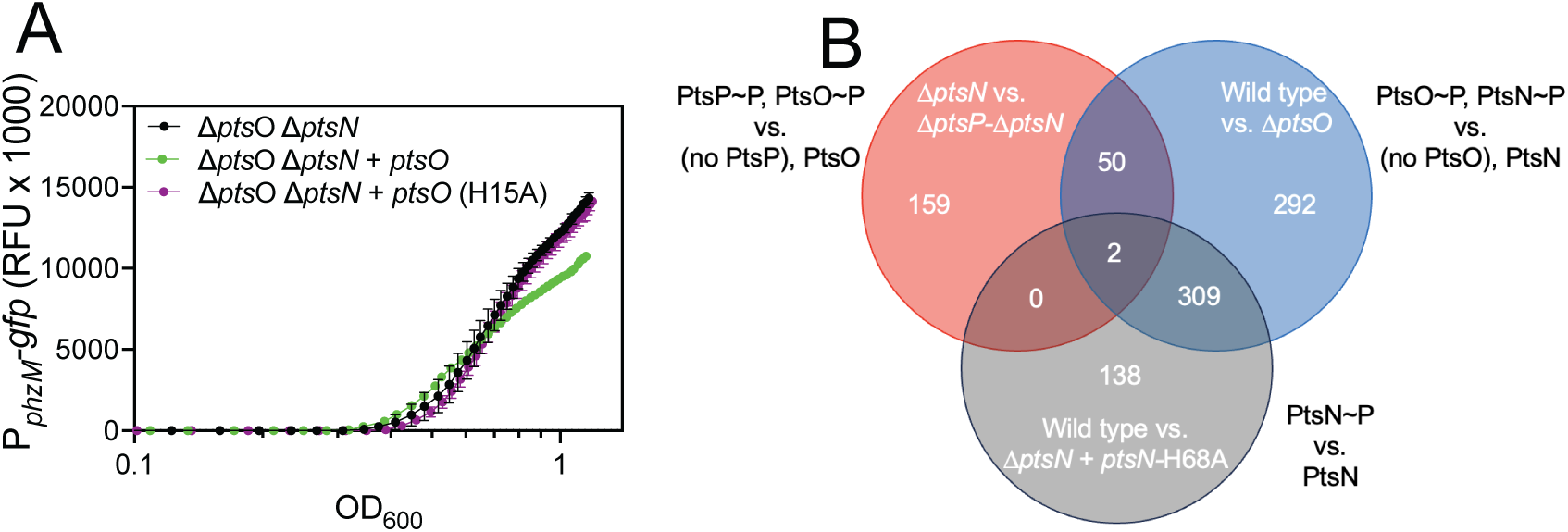
PtsO regulates a unique set of genes independent of PtsN. (A) PtsO decreases transcription from the *phzM* promoter due to its phosphorylation site. Fluorescence output measured over a time course in 96-well plates using a BioTek plate reader. (B) Venn diagram showing overlap of genes downregulated in the first strain compared with the second strain listed, as indicated, with genes >2-fold differentially expressed for each comparison. Genes for the wild type vs. Δ*ptsN* + *ptsN-*H68A were from re-analyzing data from Underhill et al. (7).

Our finding that PtsO regulates both *phzM* and *lasI* independent of the effects of PtsN led us to ask whether PtsO regulates other genes in a similar manner. To this end, we took advantage of previously conducted transcriptomic studies aimed at elucidating the PtsN regulon (7). This prior analysis included wild type, Δ*ptsN* and Δ*ptsP-*Δ*ptsN* strains, and we added an unpublished Δ*ptsO* transcriptome that was collected and analyzed with the strains from the original experiment. This combination of strains allowed us to infer whether PtsO has a distinct regulon from PtsN. We included genes regulated >2-fold to capture relatively small regulatory effects such as those we observed with *lasI*.

To determine the PtsO regulon, we first compared differentially expressed genes in wild type and Δ*ptsO* strains to identify genes impacted by the presence of PtsO (presumed to be phosphorylated in the wild-type strain) vs. no PtsO. Note that PtsN phosphorylation also differs between these strains, as PtsN is unphosphorylated in Δ*ptsO* (7). We identified 653 genes that were downregulated in the wild type compared with Δ*ptsO.* We then compared differentially expressed genes in Δ*ptsN* and Δ*ptsP-*Δ*ptsN* as another way to deduce genes regulated by PtsO; in this case the strains were presumed to have different phosphorylation states of PtsO (phosphorylated in Δ*ptsN* vs. unphosphorylated in *ΔptsP-ΔptsN).* The absence of PtsN in these strains eliminates any regulatory effects due to PtsN phosphorylation, but we note that the presence of PtsP differs between these strains. We identified 211 downregulated genes in Δ*ptsN* compared with Δ*ptsP-ΔptsN,* and more importantly, 52 genes downregulated in both the first and second strain comparisons (Fig. 6B and Table S3). Because PtsO is phosphorylated in the first strain in both comparisons, this list of 52 genes includes genes downregulated by phosphorylated PtsO.

In principle, the 52 genes identified above could also include genes downregulated by both phosphorylated PtsP and PtsN, which also differ in the first and second comparison, respectively. To this end, we included a third comparison that served to exclude genes downregulated by phosphorylated PtsN. This third comparison was of wild type and Δ*ptsN* complemented with the PtsN^H68A^ phosphorylation site mutant. In this comparison, we uncovered 449 genes downregulated by phosphorylated PtsN (Fig. 6B), with a majority (69%) of these genes also downregulated in the wild type vs. Δ*ptsO* comparison, presumably due to the difference in PtsN phosphorylation, which is common to both comparisons. Importantly, 50 of the 52 genes from our first two strain comparisons were excluded from those identified to be regulated by PtsN. These results lend confidence in these 50 genes as candidates for repression by phosphorylated PtsO.

Among the 50 candidate PtsO-repressed genes, the most highly regulated were those encoding tRNAs (tRNA-Leu, tRNA-Glu and tRNA-Gly), 5S rRNA, and the aminoglycoside-specific MexXY efflux pump. Notably, the *mexXY* genes were more strongly downregulated in Δ*ptsN* vs. Δ*ptsP*-Δ*ptsN* (∼45-fold) than in the wild type compared with Δ*ptsO* comparison (∼3-fold), suggesting these genes may have differing responses to different PTS^Ntr^ enzymes. Of note, neither *lasI* nor *phzM* were among this list of 50 genes; however, their absence might be explained by differences in growth conditions (lysogeny broth for reporter experiments vs. synthetic cystic fibrosis sputum medium for transcriptomic studies). Nevertheless, the results of our transcriptomic analysis support the idea that PtsO and PtsN can have independent effects on gene regulation.

## Discussion

In this study, we examined the regulatory link between PTS^Ntr^ and quorum sensing in *P. aeruginosa*. We identified several quorum sensing-regulated genes that are also regulated by PTS^Ntr^. By examining regulatory effects on *lasI,* the gene encoding the synthase of the LasR-specific signal, we delineated the role of each of the three PTS^Ntr^ enzymes in gene regulation and showed that modulating phosphorylation of either PtsO or PtsN is sufficient to impact *lasI* gene expression. Our results suggest a model where blocking phosphotransfer through the Nitro-PTS^Ntr^ mediates PtsN-dependent effects through unphosphorylated PtsN and PtsN-independent effects through PtsO (Fig. 7). These data support the notion that the constituent enzymes of PTS^Ntr^ have roles beyond mere phosphotransfer to PtsN. Consistent with this idea, we also identified a PtsO-specific regulon that is distinct from that of PtsN.

**Fig. 7.**
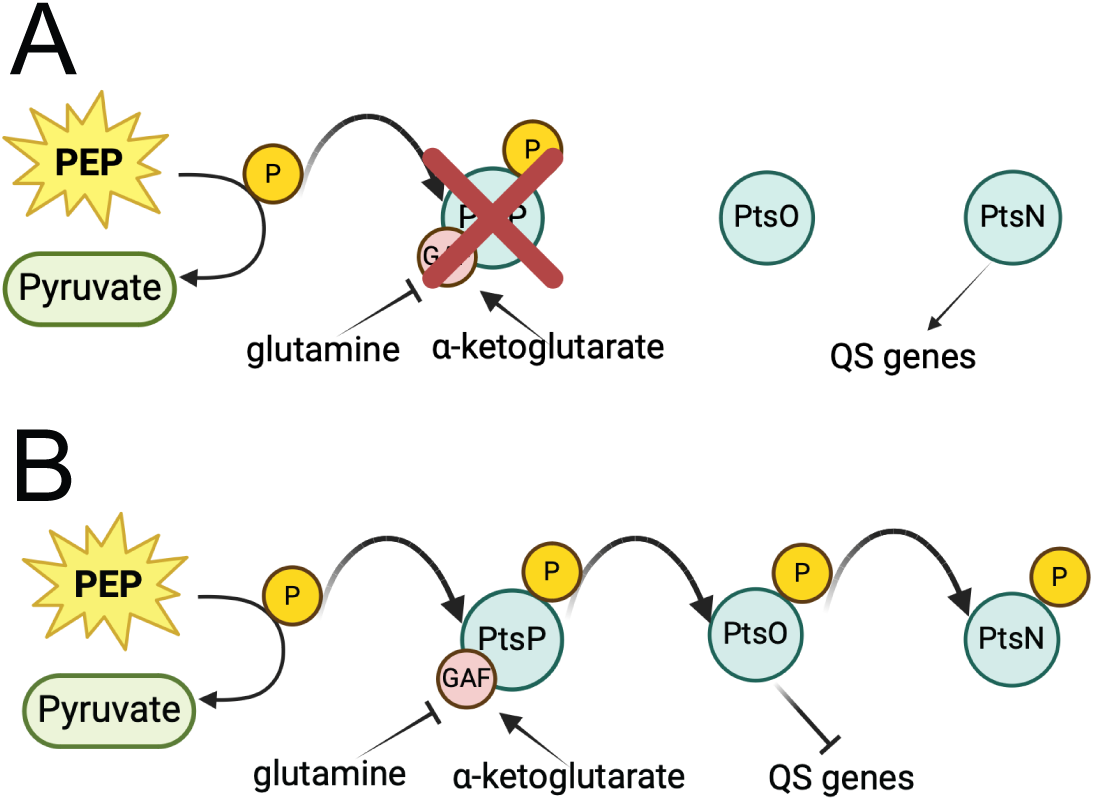
Model of PTS^Ntr^ regulation. (A) When PtsP is blocked, PtsO and PtsN is unphosphorylated, and unphosphorylated PTsN activates quorum sensing genes (e.g. *lasI*). (B) When PtsP is not blocked there is phosphate flow leading to phosphorylation of PtsO and PtsN, and phosphorylated PtsO represses quorum sensing genes.

Collectively, our study adds to the growing body of knowledge of the PTS^Ntr^ in *P. aeruginosa* and provides new information on its connection to quorum sensing regulation. In the absence of PtsP, additive effects of both PtsN-dependent and PtsN-independent regulation increase *lasI* expression (Figs. 2A, 3C). The increase in *lasI* expression causes a corollary increase in the LasI-dependent quorum sensing signal, 3OC12-HSL (Fig. 2), although this increase in 3OC12-HSL is not sufficient to activate quorum sensing more generally, as not all the LasR-LasI-dependent genes are transcriptionally activated in a Δ*ptsP* mutant (e.g. *lasB;* Fig. 1). The mechanism by which the PTS^Ntr^ interacts with the Las quorum-sensing system warrants further inquiry.

A major finding of our study is that PtsO has distinct regulatory effects that are independent of its role in phosphotransfer to PtsN. Evidence from other species corroborates this finding: for example, PtsO regulates lipid A biosynthesis (35, 36) and envelope stress responses (37) in *E. coli.* The conservation of PtsN-independent regulation by PtsO across other species suggests there may be a benefit of PtsO and PtsN acting as distinct regulatory outputs of PTS^Ntr^. One possible benefit is to enable a greater range or sensitivity to different inputs. For example, PtsN can be phosphorylated by the PTS^Fru^ enzyme FruB in both *P. aeruginosa* (7) and *P. putida* (38). In principle, FruB phosphorylation of PtsN might lead to differential phosphorylation states of PtsN and PtsO. This type of regulation might enable the cells to respond independently to changes in fructose concentrations and to other inputs of the PTS^Ntr^ system.

In our transcriptomic analyses, the presence of phosphorylated PtsO was associated with suppressed synthesis of tRNAs that can be charged with lysine, leucine, selenocysteine, glutamate and glycine. These results suggest phosphorylated PtsO might alter synthesis of certain proteins based on their amino acid composition. The depletion of certain tRNAs could also slow down translation or cause ribosome stalling, which could serve to induce certain stress responses (e.g., the stringent response). It is notable that the *mexXY* genes, which are known to be induced by ribosome stalling (39–41), were also downregulated by phosphorylated PtsO, which could be an indirect effect from the effects on translation. These genes were also more strongly repressed in the Δ*ptsN* vs. Δ*ptsP ΔptsN* comparison than in the wild type vs. Δ*ptsO* comparison suggesting *mexXY* may have an additional level of repression by PtsP, which differs in the first comparison and not in the second. Further studies of the link between the PTS^Ntr^ and MexXY, antibiotic resistance, or both, are needed to better understand this connection.

To date, few direct PTS^Ntr^ targets have been identified. The protein targets of the PTS^Ntr^ phosphoenzymes are assumed to be regulated through protein-protein interactions. In *E. coli,* TrkA, a K^+^ transporter, was identified as the direct target of PtsN (42), and the *P. putida* PtsN directly targets the PDH enzyme complex, which converts pyruvate to acetyl-CoA in the TCA cycle (43, 44). To our knowledge, no direct targets have been studied in *P. aeruginosa.* Nonetheless, our studies of quorum sensing support the idea that PtsO and PtsN may have unique direct targets that each regulate different downstream genes.

## MATERIALS AND METHODS

### Bacterial culture conditions and reagents

Bacteria were routinely grown in Lysogeny Broth (LB) (if *Escherichia coli*) or LB buffered to pH 7 with 50 mM 3-(morpholino)-propanesulfonic acid (MOPS) (if *Pseudomonas aeruginosa*), or on LB agar (LBA; 1.5% w/v Bacto-Agar; for both *E. coli* and *P. aeruginosa*). All *P. aeruginosa* broth cultures were grown in 18 mm test tubes (for 2 ml cultures) at 37°C with shaking at 250 rpm, 125 ml baffled flasks (for 10 ml cultures), or 250 ml baffled flasks (for 50 ml cultures), unless otherwise specified. For *E. coli*, 10-20 µg ml^-1^ gentamicin (depending on the strain), 10 µg ml^-1^ tetracycline and 100 µg ml^-1^ ampicillin were used. For *P. aeruginosa*, 50–200 µg ml^-1^ gentamicin (on LBA), 15 µg ml^-1^ gentamicin (in LB broth), and 200 µg ml^-1^ tetracycline were used. The CTX-2-P_ara_-*lasI* strains were grown in either 0% or 0.5% arabinose to induce plasmid expression. N-3-oxo-dodecanoyl-L-homoserine lactone (3OC12-HSL) was purchased from Cayman Chemicals (Ann Arbor, MI, USA), dissolved in acidified ethyl acetate (ethyl acetate mixed with 0.1 ml l^−1^glacial acetic acid) (45), and added to culture tubes and dried using nitrogen gas prior to adding cultures. Genomic or plasmid DNA was extracted using Qiagen Puregene Core A kit (Hilden, Germany) or IBI Scientific plasmid purification mini-prep kit (IA, USA) while PCR products were purified using IBI Scientific PCR clean-up/gel extraction kits, according to the manufacturer’s protocol. Gentamicin antibiotics was purchased from GoldBio (MO, USA), tetracycline was purchased from Fisher Scientific (PA, USA), and ampicillin was purchased from Sigma Aldrich (MO, USA). To measure luminescence (β-galatosidase) activity, the Galacto-Light Reaction Tropix kit from ThermoFisher Scientific (PA, USA) was used.

### Bacterial strains and strain construction

All bacterial strains, plasmids, and primers used in this study are listed in Tables 1-3. *P*. *aeruginosa* strain UCBPP-PA14 (‘PA14’)(46) and PA14 derivatives were used for these studies. Allelic exchange was used to make markerless deletions in specific loci of *P*. *aeruginosa* PA14 as described elsewhere (47). For allelic exchange, DNA fragments carrying the mutated or deleted gene allele plus 500 bp flanking DNA were synthesized by GenScript and inserted into the pEXG2 suicide vector. During the process of allelic exchange, the plasmids were moved to *P. aeruginosa* by conjugation using an *E. coli* donor strain and transformants were selected on Pseudomonas Isolation Agar (PIA) using gentamicin (200 μg ml^-1^) and counterselected on NaCl-free LB agar containing 15% sucrose. Putative mutants were verified through antibiotic sensitivity tests and gene-targeted Sanger sequencing. CTX plasmids were also moved to *P. aeruginosa* strains via conjugation from an *E. coli* donor strain. DNA fragments from CTX that incorporated at the neutral chromosomal *attB* locus were selected on PIA-Tetracycline (200 μg ml^-1^) and verified by PCR (48, 49). All replicating plasmids were introduced to *P. aeruginosa* via electroporation (30), selected on LB agar using gentamicin at 50–200 μg ml^-1^ and routinely grown with gentamicin (50 μg ml^-1^ for agar and 15 μg ml^-1^ for broth) for plasmid maintenance.

### Reporter activity measurements

The influence of the PTS^Ntr^ system on *lasI, phzM, rhlA, lasB, rsaL,* and *hcnA* gene expression levels in *P. aeruginosa* was measured using transcriptional reporter plasmids with each gene promoter fused to the promoterless *gfp* reporter in the pPROBE plasmid. Each plasmid was moved to *P. aeruginosa* by electroporation, and single transformant colonies were used to inoculate 2 mL LB-MOPS Gm_15_ to start the experiments. For end-point measurements of activity, cells were grown for 18 h, washed with phosphate buffered saline (PBS), suspended in PBS, and fluorescence and optical density were measured using a BioTek Synergy 2 plate reader. For time-course measurements, overnight cultures were diluted 1:100 in fresh medium and grown to an optical density at 600 nm (OD_600_) of ∼0.1. This culture was then diluted to an OD_600_ of ∼0.004 and dispensed in a black 96-well clear flat-bottom plate with 200 µl per well. Plates were incubated with double orbital shaking at 37 °C in a Biotek Synergy H1 plate reader with OD_600_ and GFP fluorescence (excitation 485 nm, emission 528 nm) measured every 15 min for 14 h. To account for background, the empty vector pPROBE P*_empty_*-gfp was also measured and subtracted from the measured values of each reporter strain. The final fluorescence values were plotted with respect to OD_600_ using GraphPad Prism.

### Quorum sensing signal measurements

We used a bioassay to measure 3OC12-HSL produced by different strains of *P. aeruginosa.* To prepare samples for analysis, we used 5 ml cultures grown 18 h in LB-MOPS. The cells were removed from the culture fluid by centrifugation, the culture fluid was extracted twice with acidified ethyl acetate, and the ethyl acetate fraction was evaporated to dryness under a stream of nitrogen gas. The residue was dissolved in 0.5 ml acidified ethyl acetate, and the ethyl acetate solutions were used in bioassay. For the bioassay, we used strain *E. coli* DH5a pSC11 pJ105L-LasR with P*lasI-lacZ* and *lasR* on different plasmids (30). Details of the bioassay procedure have been described elsewhere. Briefly, overnight *E. coli* cultures grown in LB-MOPS were diluted 1:100 into fresh LB-MOPS and grown to an OD_600_ of ∼0.2-0.3 prior to adding arabinose for induction of *lasR.* The culture was then grown to an OD_600_ of ∼0.5-0.6, and aliquots (0.5 ml) were dispensed into 2 ml Eppendorf tubes with evaporated culture fluid extracts, synthetic 3OC12-HSL standards or no signal and grown an additional 3 h. β-galactosidase was detected using the Galacto-light Reaction Tropix Kit (Thermo Fisher) and measured using a Biotek Synergy 2 plate reader. A standard curve was generated from the synthetic signal samples and signal levels in the culture fluid extracts were determined by comparing with the standard curve.

### RNA extraction and sequencing

RNA extraction and sequencing was as described previously (7). Briefly, cultures grown to an OD600 of 0.3 in SCFM2 were harvested and RNA was extracted according to the NEB (New England Biolabs, MA, USA) Monarch RNA extraction kit. rRNA was depleted using the Illumina RiboZero kit (Illumina, CA, USA) and samples were sequenced by 150-bp paired-end Illumina sequencing at the University of Oklahoma Health Sciences Center core facility in Oklahoma City, OK, USA. Sequence mapping and analysis were performed at the Oklahoma University Health Sciences Center Laboratory for Molecular Biology and Cytometry Research using CLC software. Genes were classified as differentially expressed in pair-wise comparisons based on the FDR-adjusted P-value < 0.001 and log_2_ fold-change values > 1, and a summary of the overlap in differentially expressed genes between pair-wise comparisons was generated with the R package ggVennDiagram v 1.5.2 in R v 4.4.1 (50, 51).

## ACKNOWLEDGEMENTS

This work was supported by the NIH through grants R35GM156409 to JRC, R35GM138018 to MTC, and R35GM152107 to AAD, as well as the Oklahoma Center for Microbial Pathogenesis and Immunity (GM134973). The transcriptomics re-analysis was carried out by Dr. Brian Sanderson and supported by K-INBRE, which is supported by the NIGMS IDeA Program award number P20 GM103418, as well as the University of Kansas Center for Genomics.

