## Supplemental material for "Quorum sensing regulation by the nitrogen phosphotransferase system in *Pseudomonas aeruginosa*"

Supplementary methods

**Western blot.** Cells for Western blot were grown in test tubes at 37°C with shaking for 18hrs. At 18hrs, at which point the cell densities were measured at an optical density of 600 nm ($OD_{600}$) and determined to be equivalent within 5%. Samples (1 ml) were then centrifuged, and the cell pellets were washed twice with cold 1X TBS (2 0mM Tris-HCl, pH 7.5, 150 mM NaCl), and the washed cell pellets were subjected to a cycle of freeze-thaw at -80°C. The cell pellet was then dissolved in 50 mM Tris buffer, using 50 µL of Tris buffer (50 mM Tris-HCl, pH 7.5), and lysed by adding 100 µL loading buffer (60 mM Tris pH 6.8, 2% sodium dodecyl sulfate [SDS], 10% Glycerol, 0.01% bromophenol blue) supplemented with 1X Protease Inhibitor Complex (PIC) and Phosphatase Inhibitor Complex (PhIC), 1% Phenylmethylsulfonyl fluoride (PMSF) and 1% β-Mercaptoethanol (BME) and this was treated with 1 µl Universal Nuclease for 1hr at 37°C. The samples were heated at 95°C for 5-10 minutes, centrifuged at 16,000g for 20mins and supernatant was collected. The supernatant was further diluted by sample buffer (1:5 ratio) in 2X SDS sample buffer and equal amounts of sample for each strain was loaded. For Western blots, primary antibody (DYKDDDDK Tag Antibody, mAb, Mouse, from Genscript) and secondary antibody (Goat anti-mouse, IRDye 680RD, from LICOR) was used and membranes were imaged using a LICOR Odyssey M imager.

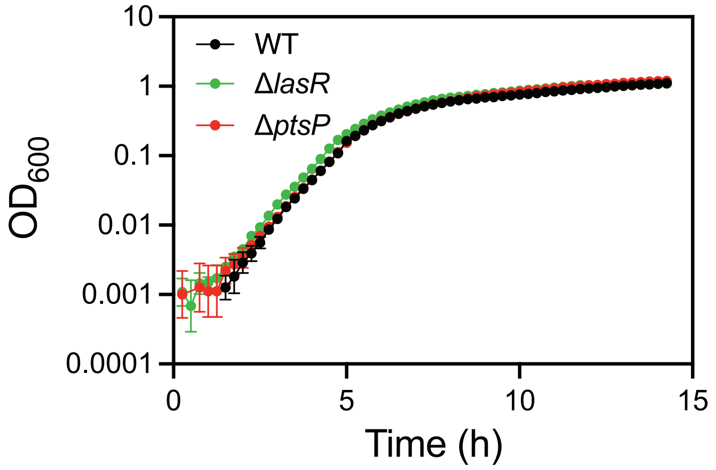


**Figure S1. Growth of strains in experiments presented in Fig. 1.** Data show growth of PA14 strains transformed with the pP$_{lasI}$-*gfp* plasmid; all other reporter plasmids showed similar results. Data points are the means of three replicates, and the error bars represent standard deviation. All strains were grown in MOPS-buffered LB.

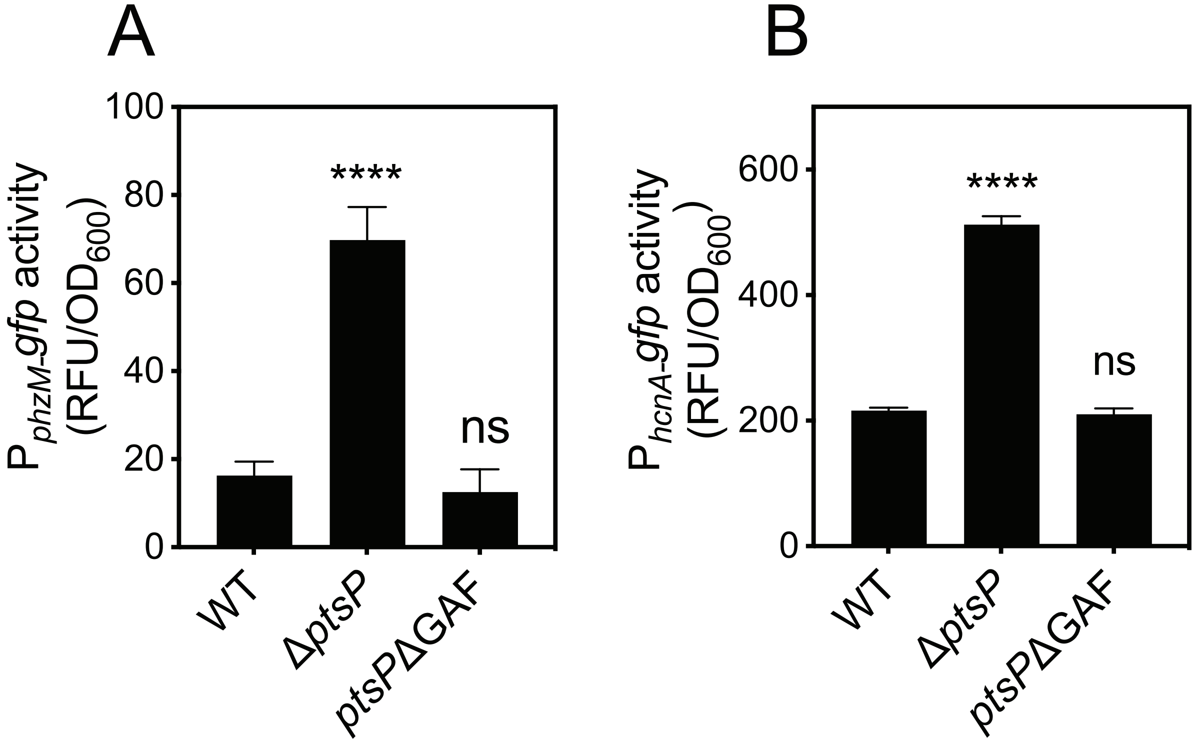


**Figure S2. The GAF domain of PtsP is indepensible for its role in regulating transcription from the *phzM* (A) and *hcnA* (B) promoters.** Transcription was monitored as GFP fluorescence in cells transformed with the $pP_{phzM}$-*gfp* or $pP_{hcnA}$-*gfp* reporter plasmids. Data shown are growth-adjusted fluorescence after 16 h growth.. Data are means of at least three replicates, and error bars represent SD. Statistical significance by one-way ANOVA compared with wild type; ****, $p < 0.0001$; ns, not significant.

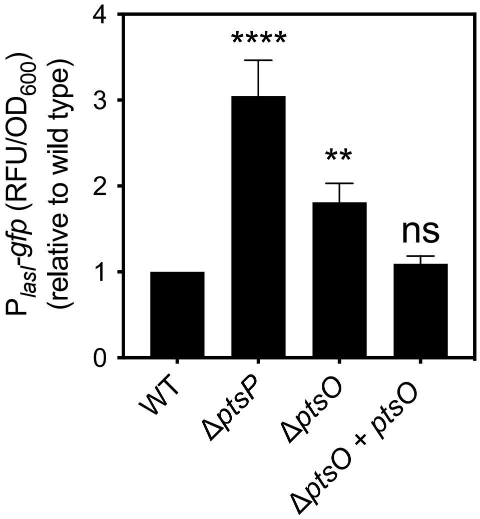


**Figure S3. Δ*ptsO* complementation.** Transcription from the *lasI* promoter was monitored as GFP fluorescence in cells transformed with the p$P_{lasI}$-*gfp* reporter plasmid. Strains carried *ptsO* or the empty CTX cassette in the neutral *attB* site in the genome. Fluorescence was obtained after overnight growth and is normalized to $OD_{600}$ and shown as the fold change compared with wild type. Data are means of at least three replicates, and error bars represent SD. Statistical significance by one-way ANOVA compared with wild type; ****, $p < 0.0001$; **, $p < 0.005$; ns, not significant.

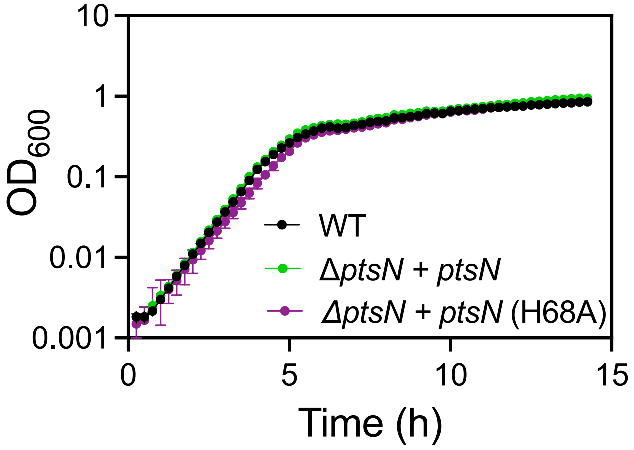


**Figure S4. Growth of *ptsN* mutants.** Growth curves for experiments presented in Fig. 4B in the main text. As in Fig. 4B, strains carried the pP$_{lasI}$-*gfp* reporter plasmid and *ptsN, ptsN* (H68A) or an empty cassette inserted at the neutral *attB* site in the chromosome and measurements were taken on a Biotek plate reader. Data points are the means of three replicates, and the error bars represent standard deviation. All strains were grown in MOPS-buffered LB.

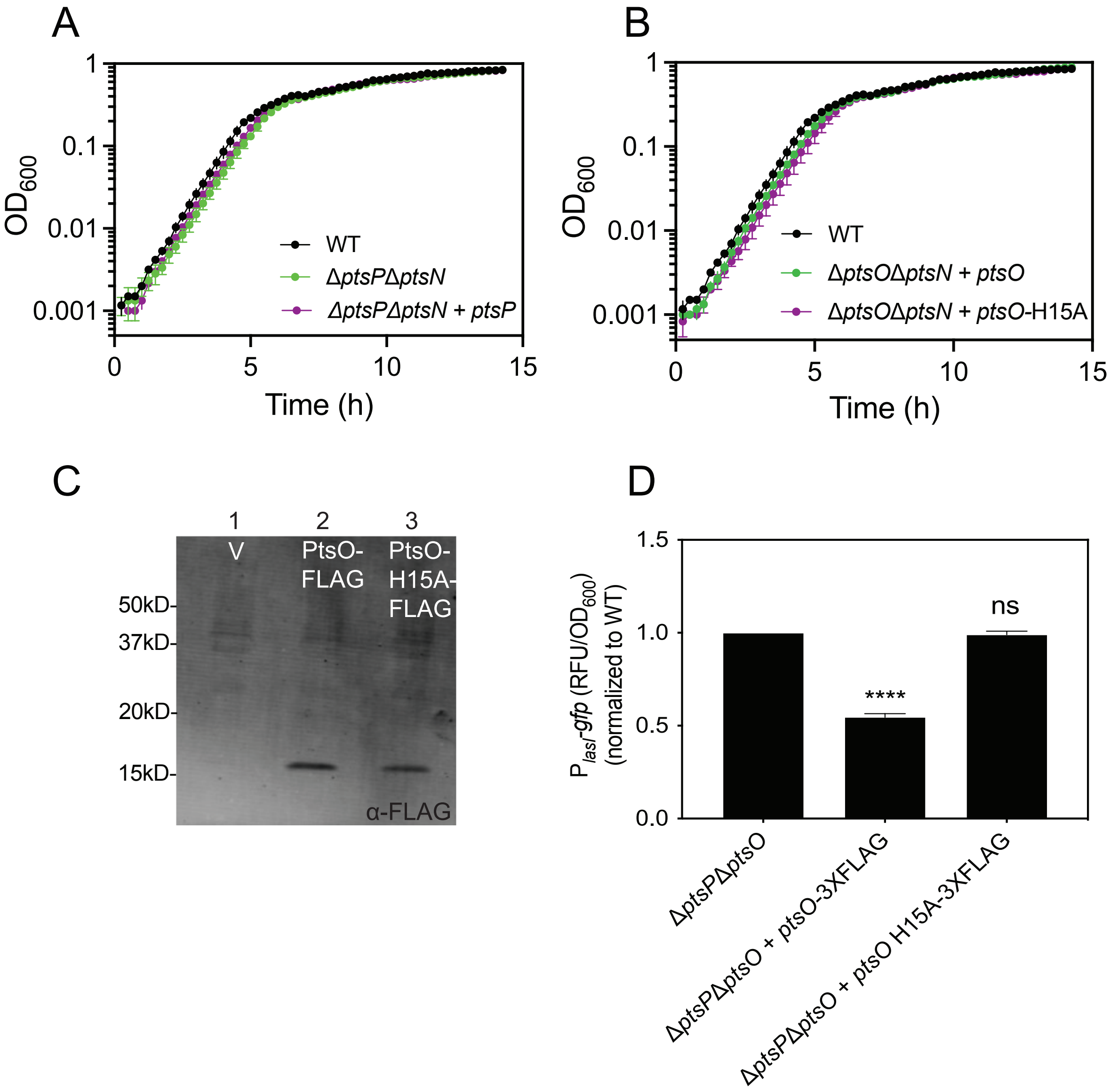


**Fig. S5. Effects of PtsO and PtsO H15A on growth and PtsO levels.** (A) Growth of strains in experiments presented in Figs. 5B (A) and 5D (B). In both panels, all strains were carrying the pP$_{lasI}$-*gfp* plasmid and grown in MOPS-buffered LB. Data points are the means of three replicates, and the error bars represent standard deviation. (C) Western blot to monitor levels of PtsO protein in whole cell lysates. Strains are Δ*ptsO*-Δ*ptsN* with 3X-FLAG-tagged *ptsO* or *ptsO* H15A introduced by the CTX integration vector. V (lane 1), Δ*ptsO*-Δ*ptsN* carrying the empty CTX vector only. Strains were grown to stationary phase (18 h) in MOPS-buffered LB prior to harvesting and were at equivalent cell densities as measured by the optical density at 600 nm ($OD_{600}$). (D) To demonstrate that addition of 3XFLAG tag does not cause any unknown effects on PtsO repression of *lasI* transcription, we used the pP*lasI*-*gfp* reporter plasmid to monitor *lasI* expression at 18 h of strains grown as in panel C. Results are normalized to $OD_{600}$ and shown as the fold change compared with wild type. Data are means of at least three replicates, and error bars represent SD. Statistical significance by one-way ANOVA compared with Δ*ptsO*-Δ*ptsN*; ****, $p < 0.0001$; ** ; ns, not significant.

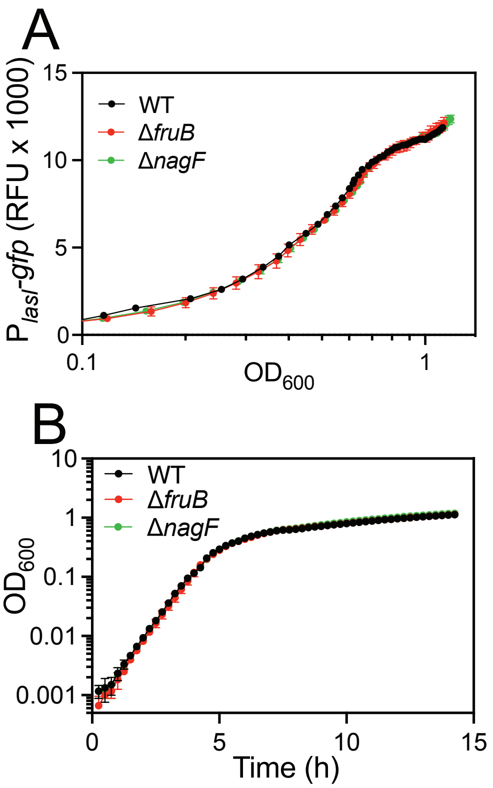


**Fig. S6. *lasI* expression levels in strains with deletions of the carbohydrate phosphotransferase genes *nagF* and *fruB.*** (A) Transcription from the *lasI* promoter was monitored as GFP fluorescence in cells transformed with the $pP_{lasI}$-*gfp* reporter plasmid. Fluorescence output measured over a time course in 96-well plates using a BioTek plate reader. (B) Growth of strains in experiments presented in panel A. Data points are the means of three replicates, and the error bars represent standard deviation.

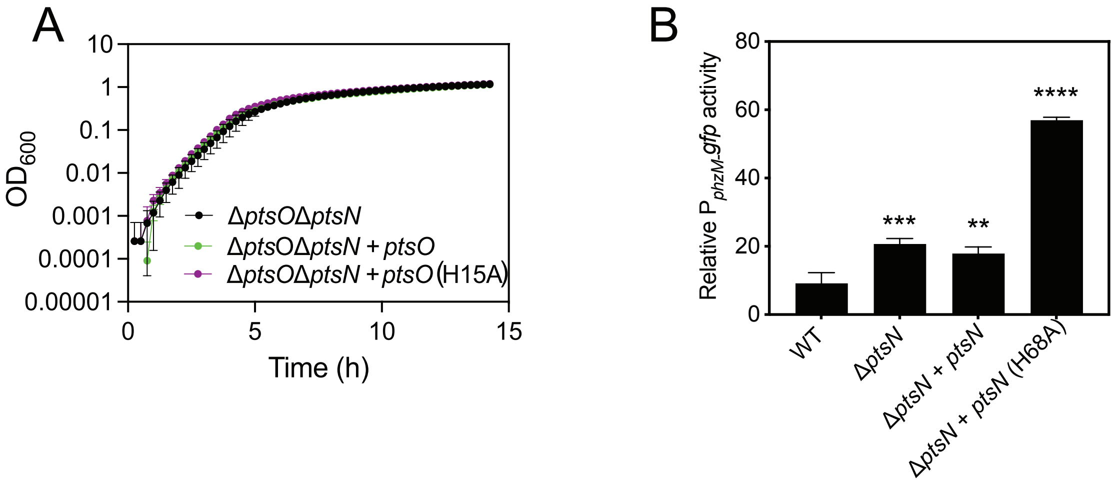


**Fig. S7. Regulation of *phzM* expression by PtsN and PtsO.** (A) Growth of strains in experiment presented in Fig. 6A. In both panels, all strains were carrying the pP$_{phzM}$-*gfp* plasmid and grown in MOPS-buffered LB. Data points are the means of three replicates, and the error bars represent standard deviation. Transcription from these promoters was monitored as GFP fluorescence in cells transformed with plasmid-based reporters. (A) Strains carried a chromosomally integrated CTX-1 cassette (CTX), CTX plus the wild type *ptsO,* or CTX plus the H15A variant of PtsO (*ptsO* (H15A)). Fluorescence output was measured over a time course in 96-well plates using a BioTek plate reader. (B) Strains carried CTX, CTX plus the wild-type *ptsN,* or CTX plus the H68A variant of PtsN (*ptsN* (H68A)). Fluorescence was obtained after overnight growth and is normalized to $OD_{600}$. Data are means of at least three replicates, and error bars represent SD. Statistical significance by one-way ANOVA compared with Δ*ptsP*-Δ*ptsN* + *ptsN*-H68A; ****, $p < 0.0001$; **, $p < 0.005$; ns, not significant.

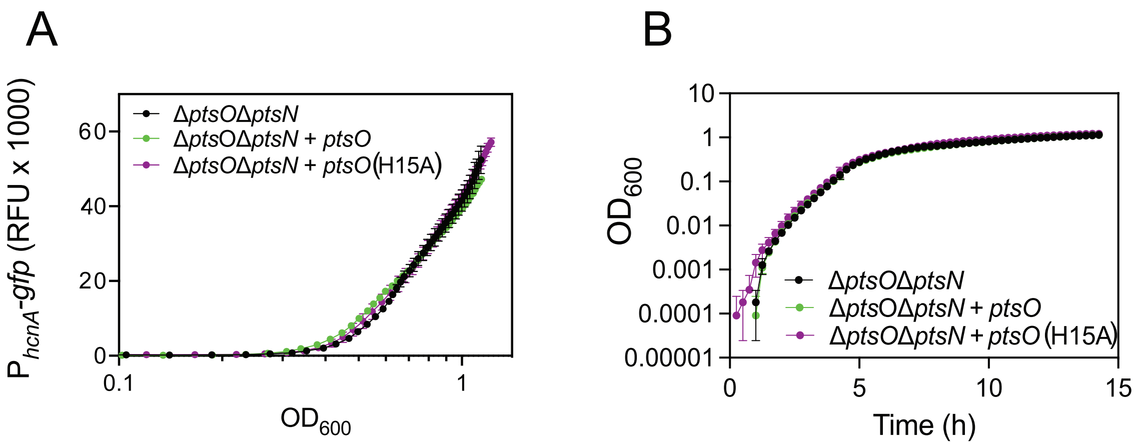


**Fig. S8. *hcnA* expression is not modulated by PtsO.** (A) Transcription from the *hcnA* promoter was monitored as GFP fluorescence in cells transformed with the pP$_{hcnA}$-*gfp* reporter plasmid. Fluorescence output was measured over a time course in 96-well plates using a BioTek plate reader. Strains carried a chromosomally integrated CTX-1 cassette (CTX), CTX plus the wild type *ptsO,* or CTX plus the H15A variant of PtsO (*ptsO* (H15A)). (B) Growth of strains in experiments presented in panel A. Data points are the means of three replicates, and the error bars represent standard deviation.

Table S1. Bacterial strains used in this study.

| Strains | Relevant properties | Reference or source |
|---|---|---|
| *P. aeruginosa* strains | | |
| UCBPP- PA14 (PA14) | Ancestral wild type | (1) |
| PA14 Δ*ptsP* | PA14 with a deletion of *ptsP* | (2) |
| PA14 Δ*lasR* | PA14 with a deletion of *lasR* | This study |
| PA14 Δ*lasI* | PA14 with a deletion of *lasI* | (2) |
| PA14 Δ*ptsO* | PA14 with a deletion of *ptsO* | (3) |
| PA14 Δ*ptsN* | PA14 with a deletion of *ptsN* | (3) |
| PA14 Δ*ptsP*-Δ*ptsO* | PA14 Δ*ptsO* with a deletion of *ptsP* | This study |
| PA14 Δ*ptsP*-Δ*ptsN* | PA14 Δ*ptsN* with a deletion of *ptsP* | This study |
| PA14 Δ*ptsP*-Δ*ptsO*-Δ*ptsN* | PA14Δ*ptsP*Δ*ptsN* with a deletion of *ptsO* | This study |
| PA14 Δ*lasI*-Δ*ptsP* | PA14 Δ*lasI* with a deletion of *ptsP* | This study |
| PA14 *ptsP*ΔGAF | PA14 in which the coding sequence of the PtsP GAF domain has been deleted | (3) |
| PA14 Δ*ptsO*-Δ*ptsN* | PA14 with *ptsO* and *ptsN* gene deletions | (4) |
| *Escherichia coli* strains | | |
| E.coli DH5α | $F^-$ ϕ80*lacZ*ΔM15 Δ(*lacZYA-argF*)*U169 hsdR17*($r_K^-$ $m_K^+$) *recA1 endA1 phoA supE44 thi-1 gyrA96 relA1* $\lambda^-$ | Invitrogen |
| E.coli S17-1 | *recA pro hsdR* RP4-2-Tc::Mu-km::Tn*7* | (5) |
| E.coli SM10 | *thi thr leu tonA lacY supE recA*::RP4-2-Tc::Mu Km λ*pir* | (5) |
| E.coli DH5α pSC11 pJ105L-LasR | 3OC12-HSL-responsive bioassay strain | (6) |

Table S2. Plasmids used in this study.

| Strains | Relevant properties | Reference or source |
|---|---|---|
| pPROBE-GT | Broad-host-range pVS1/p15a GFP reporter; $Gm^R$ | (7) |
| pCTX-1 | Mini-CTX, *P. aeruginosa* integrative plasmid; $Tet^R$ | (8) |
| pCTX-2 | Mini-CTX with arabinose-inducible promoter; $Tet^R$ | (9) |
| pEXG2 | Suicide plasmid for allelic replacement; $Gm^R$ | (10) |
| pBS351:$P_{lasI}$-*gfp* | Encodes −1 through −501 relatives to the start of *lasI* | (6) |
| pPROBE-$P_{rsaL}$-*gfp* | Encodes +103 to −290 relatives to the start of *rsaL* | (11) |
| pPROBE-$P_{phzM}$-*gfp* | Encodes +20 to −350 relatives to the start of *phzM* | This study |
| pPROBE-$P_{rhlA}$ -*gfp* | Encodes +1 through −501 relatives to the start of *rhlA* | (6) |
| pPROBE $P_{lasB}$ -*gfp* | Encodes +217 to −290 relatives to the start of *lasB* | (12) |
| pPROBE $P_{hcnA}$ -*gfp* | Encodes +1 through −500 relatives to the start of *hcnA* | This study |
| pCTX1-*ptsP* | CTX-1 containing the *ptsP* gene, $Tet^r$ | (3) |
| pSW196-RBS-*lasI* | Mini-CTX2 with $P_{ara}$*BAD* promoter; $Tet^r$ | (13) |
| pCTX-1-Pop-*ptsN* | CTX-1 with *ptsN* fused to the *rpoN* operon promoter | (4) |
| pCTX-1-Pop-*ptsN* (H68A) | pCTX-1-Pop-*ptsN* with His 68 codon mutated to Ala | (4) |
| pCTX1-*ptsO* | PA14 *ptsO* on pCTX1 vector | This study |
| pCTX1-*ptsO* (H15A) | pCTX1-*ptsO* with His 15 codon mutated to Ala | This study |
| pCTX1-*ptsO* (3XFlag) | pCTX1-*ptsO* with 3XFlag tag at the 3' end | This study |
| pCTX1-*ptsO* (H15A-3XFlag) | CTX1-*ptsO* H15A with 3XFlag tag at the 3' end | This study |
| pEXG2 Δ*ptsO* | Used to make in-frame *ptsO* deletion | (3) |
| pEXG2 Δ*ptsN* | Used to make in-frame *ptsN* deletion | (3) |
| pEXG2 -PA14 Δ*lasR* | Used to make in-frame *lasR* deletion with *rsaL* intact | Kostylev, unpublished |
| pEXG2 Δ*fruB* | Used to make in-frame *fruB* deletion | (4) |
| pEXG2 Δ*nagF* | Used to make in-frame *nagF* deletion | (4) |
